# Ribosome provisioning activates a bistable switch coupled to fast exit from stationary phase

**DOI:** 10.1101/244129

**Authors:** P. Remigi, G.C. Ferguson, S. De Monte, P.B. Rainey

## Abstract

Observations of bacteria at the single-cell level have revealed many instances of phenotypic heterogeneity within otherwise clonal populations, but the selective causes, molecular bases and broader ecological relevance remain poorly understood. In an earlier experiment in which the bacterium *Pseudomonas fluorescens* SBW25 was propagated under a selective regime that mimicked the host immune response, a genotype evolved that stochastically switched between capsulation states. The genetic cause was a mutation in *carB* that decreased the pyrimidine pool (and growth rate), lowering the activation threshold of a pre-existing but hitherto unrecognised phenotypic switch. Genetic components surrounding bifurcation of UTP flux towards DNA/RNA or UDP-glucose (a precursor of colanic acid forming the capsules) were implicated as key components. Extending these molecular analyses – and based on a combination of genetics, transcriptomics, biochemistry and mathematical modelling – we show that pyrimidine limitation triggers an increase in ribosome biosynthesis and that switching is caused by competition between ribosomes and CsrA/RsmA proteins for the mRNA transcript of a feed-forward regulator of colanic acid biosynthesis. We additionally show that in the ancestral bacterium the switch is part of a programme that determines stochastic entry into the semi-quiescent capsulated state, ensures that such cells are provisioned with excess ribosomes, and enables provisioned cells to exit rapidly from stationary phase under permissive conditions.

## Introduction

Phenotypic variation between isogenic cells growing in homogeneous environments can have adaptive consequences, allowing populations to survive unpredictable environmental changes or promoting interactions between different cell types^1,2^. Natural selection, by means of genetic mutations affecting the integration of stochastic noise within signaling pathways, can fine-tune epigenetic switches^3–7^ but the molecular details underpinning the evolution of phenotypic heterogeneity remain poorly understood.

Opportunity to study the genetic bases of the evolution of phenotypic heterogeneity arose from a selection experiment where the capacity to switch between different colony phenotypes evolved *de novo* in the bacterium *Pseudomonas fluorescens* SBW25 (ref. 8). In this experiment, bacteria were passaged through consecutive cycles comprised of single-cell bottlenecks and negative frequency-dependent selection, a regime mimicking essential features of the adaptive immune system in animals. A genotype emerged (1B4) that forms distinct opaque or translucent colonies on agar plates. At the single-cell level, this behaviour reflects an epigenetic switch characterized by the bistable production of an extracellular capsule, resulting in the coexistence of two sub-populations of capsulated (Cap^+^) or noncapsulated (Cap^−^) cells. Capsules are made of a colanic acid-like polymer, whose production arises from the activity of the *wcaJ-wzb* locus and requires the precursor UDP-glucose. A mutation in *carB* (c2020t), a gene involved in *de novo* pyrimidine biosynthesis, is responsible for heterogeneous capsule production via a decrease in intracellular pyrimidine pools^8,9^. Additionally, the switch between capsulated and uncapsulated cells was found to be active in the ancestral gentoype devoid of the *carB* mutation – the *carB* mutation having altered the threshold at which the switch is activated – and to underpin the stochastic entry of cells into a semi-quiescent state.

Here we extend earlier work and provide mechanistic understanding of how the *carB* mutation determines capsulation heterogeneity. Of central importance is evidence that ribosome biosynthesis is up-regulated upon pyrimidine limitation and that this favors translation of a positively auto-regulated activator of capsular exopolysaccharide biosynthesis that is otherwise inhibited by CsrA/Rsm proteins. The switch comprises part of a programme that facilitates stochastic entry into a semi-quiescent state and rapid exit from this state upon realisation of permissive conditions through modulation of ribosome levels.

## Results

### Capsulation is not induced by UDP-glucose depletion

Previous work analysed the switcher genotype 1B4 (ref. 8) and the link between pyrimidine limitation (caused by a defect in CarB), growth and heterogeneous expression of capsules^9^. Particular attention was given to bifurcation of UTP flux towards DNA/RNA, or UDP-glucose (a precursor of colanic acid from which capsules are synthesised). Extensive analyses showed that capsule production was tied to entry into a semi-quiescent state triggered by reduction of flux though the pyrimdine biosynthetic pathway. The primary signalling molecule was not identified, but it was hypothesised to be a product of the pyrimidine biosynthetic pathway^9^, with UDP-glucose being a prime candidate given its role in regulation of cell size and bacterial growth^10^.

To test the hypothesis that UDP-glucose underpins the switch to capsule production, a translational (GFP) reporter fused to PFLU3655 (the primary transcriptional activator of the *wcaJ-wzb* operon, see ref. 9 and below) was introduced into the chromosome of a *galU* mutant of 1B4. The *galU* mutant is unable to convert UTP to UDP-glucose and is therefore Cap^−^ (ref. 9). The proportion of cells expressing the P*pflu3655*-GFP reporter was reduced in the mutant compared to the 1B4 switching genotype (Supplementary Figure 1). This finding was inconsistent with the prediction that low UDP-glucose is the signal that increases the chance of switching to the capsulated state. Accordingly the mechanistic links between pyrimidine limitation, growth and the heterogeneous production of capsules were reassessed.

### Ribosomes are over-produced in *carB* mutants

Transcriptomic data published previously^9^ was interrogated to identify signalling pathways displaying different levels of activity as a result of the causal *carB* mutation. KEGG-enrichment analyses showed over-representation of ribosomal components among the genes that are expressed at least two-fold more in the capsulated sub-population of 1B4 (1B4 Cap^+^) compared to its (non-capsulated) ancestor 1A4 (Supplementary Table 1). By extracting raw expression values from the available RNAseq datasets, it was found that the average expression levels of ribosomal protein genes increased in 1B4 (in both Cap^−^ and Cap^+^) compared to 1A4 or SBW25 (Fig. 1a). Such a finding is surprising since ribosome production is usually proportional to growth rate^11,12^ and was expected to be reduced in the slower growing strain 1B4 (Supplementary Figure 2).

Bacteria adjust ribosome concentration to match nutrient availability in order to maximise growth rate. They do so by modulating transcriptional activity at ribosomal RNA (*rrn*) operon promoters^11,12^ causing the production of ribosomal proteins to match available rRNA (ref. 13). The over-expression of ribosomal protein genes in 1B4 may therefore reflect transcriptional up-regulation at *rrn* promoters. Using a chromosomally-integrated reporter, an increase in P*rrnB-*GFP transcriptional activity was detected in *carB* mutants compared to immediate ancestral types (Fig. 1b). This difference was most obvious when cultures reached OD~2, a density at which 1B4 cultures undergo a noticeable increase in capsulation (Supplementary Figure 3). Supplementing growth media with 2mM uracil – a treatment known to suppress capsulation^9^ – restored wild-type P*rrnB*-GFP expression in strains carrying the mutant *carB* allele (Fig. 1b). Measurement of cellular RNA content also showed higher levels in *carB* mutants (Fig. 1c). These measurements were not affected by cell size (Supplementary Figure 4) indicating a *bona fide* increase in ribosome concentration. Together, these results show that pyrimidine limitation triggers the production of an enhanced pool of ribosomes.

**Figure 1:**
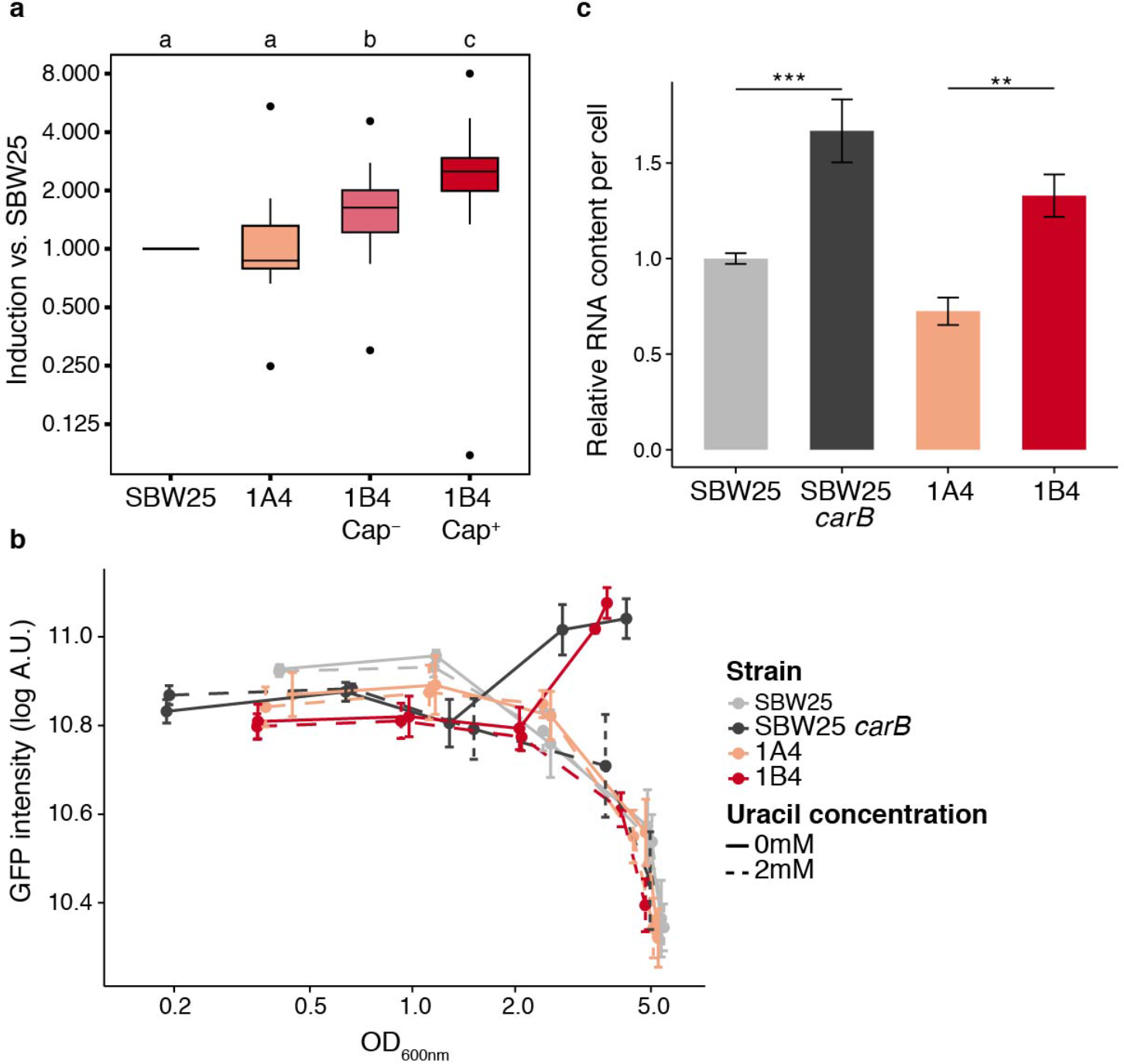
Increased ribosome production in *carB* mutants. **a**, Transcriptional induction of ribosomal protein genes in SBW25, 1A4, 1B4 Cap^−^ and 1B4 Cap^+^ cells. Absolute expression levels of ribosomal protein genes (KEGG pathway ‘0310-Ribosomes’, n=26) were extracted from a previous RNA-seq dataset^9^ and normalised to SBW25. Boxplots represent the distribution of expression ratios. Bold segments inside rectangles show the median, lower and upper limits of the box represent first and third quartiles, respectivey. Whiskers extend up to 1.5 times the interquartile range and dots represent outliers, if present. Letter groups indicate statistical significance, *P* < 0.05, Kruskall-Wallis test with Dunn’s post-hoc correction. **b**, Expression kinetics of the P*rrnB*-GFP transcriptional reporter. Fluorescence in individual cells was measured by flow cytometry. Mean fluorescence of bacterial populations ± s.d. over biological replicates are shown, n = 4. Data are representative of 3 independent experiments. **c**, Total RNA content in bacterial cells during exponential phase (OD_600nm_ = 0.5-0.6). Values were normalised to SBW25 control within each experiment. Means ± s.d. are shown, n = 6. Data are pooled from 4 independent experiments. ** *P* < 0.01, *** *P* < 0.001, two-tailed *t*-test.

### High ribosome levels are required for capsulation

The counterintuitive effect of the *carB* mutation on ribosome levels suggested a causal connection between ribosome concentration and capsulation. In support of this hypothesis, a previous transposon-mutagenesis screen found that insertions in ribosome- or translation-associated genes (*prfC, rluB, rluC, glu/gly* tRNA) decreased or abolished capsulation in 1B4 (ref. 9). We set out to manipulate ribosome concentration in 1B4 in order to test directly if ribosome abundance affects capsulation.

*P. fluorescens* SBW25 harbours five copies of *rrn* (*rrnA-E*). Because deletion of a single *rrn* operon can often be compensated by over-expression of those remaining^14–16^ capsulation was quantified in both single and double *rrn* deletion mutants using the chromosomally-inserted translational reporter P*pflu3655*-GFP (ref. 9). Whereas single mutants were not significantly affected in capsulation status, double mutants produced fewer capsulated cells (Fig. 2a and Supplementary Figure 5). Growth rate was only marginally affected in certain mutant combinations (Supplementary Figure 6) but a significant reduction in total RNA content was observed in three out of the six *rrn* double mutants (Supplementary Figure 7). Together, these results show that capsulation is positively affected by increased ribosome abundance, which is itself a response to pyrimidine starvation.

**Figure 2:**
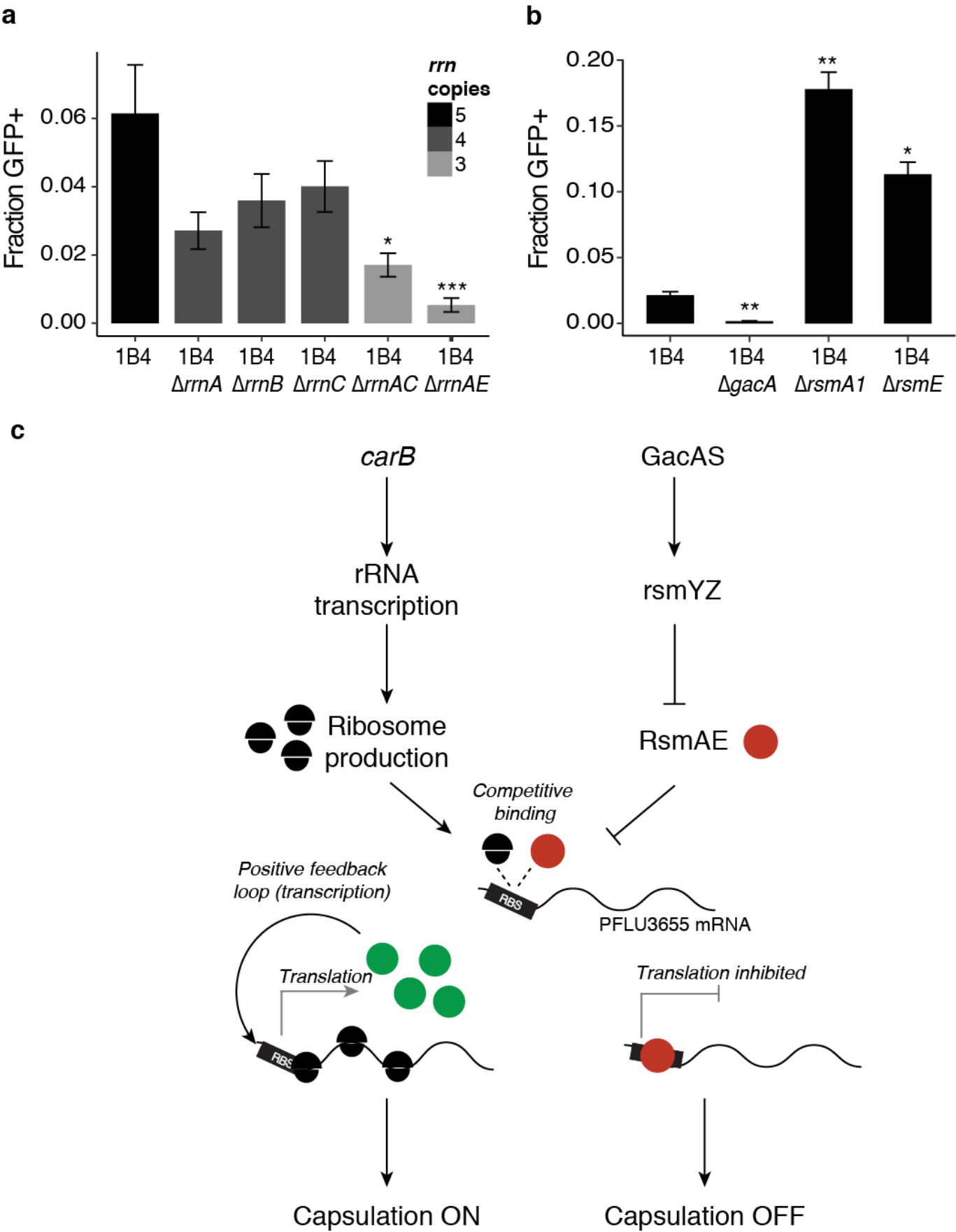
Genetic bases of capsulation. **a**, Capsulation in the rrn deletion mutants. The Tn7-P*pflu3655*-GFP reporter was introduced in 1B4 bacteria and its derived rrn mutants. Capsulation was measured by quantifying the proportion of GFP positive cells by flow cytometry at the onset of stationary phase (OD_600nm_ = 1-2). Means ± s.e.m. are shown, n = 8 (1B4 ∆*rrnB*) or n = 11 (all other strains). Data are pooled from 4 independent experiments. ** *P* < 0.01, *** *P* < 0.001, Kruskall-Wallis test with Dunn’s post-hoc correction, comparison to 1B4. **b**, Capsulation in *gac*/*rsm* mutants. Means ± s.e.m. are shown, n = 12 (1B4), n = 15 (1B4 ∆*gacA*) or n = 9 (all other strains). Data are pooled from 3 independent experiments. * *P* < 0.05, *** *P* < 0.001, Kruskall-Wallis test with Dunn’s post-hoc correction, comparison to 1B4. **c**, A model for capsulation in 1B4. See text for details.

### A ribosome-Rsm competition model for the control of capsulation

Next, we asked how ribosome abundance influences heterogeneous production of capsulated cells. Results from a previous transposon mutagenesis screen showed that the Gac/Rsm two-component signalling pathway is required for production of colanic acid-like capsules^9^. The Gac/Rsm signalling pathway controls important ecological traits in many Gram-negative bacteria, including secretion of exoproducts, the transition between biofilm and planktonic cells and pathogenicity^17,18^. Its activity is mediated through post-transcriptional regulators of the Csr/Rsm family that prevent translation of mRNA targets by binding to sites adjacent to or overlapping ribosome-binding sites (RBS)^17,19^. Upon perception of unknown extracellular signals^18^, the sensor kinase GacS activates the cognate response regulator GacA and the transcription of small non-coding RNAs *rsmY* and *rsmZ*. Binding of these sRNAs to Csr/Rsm proteins antagonizes their translation inhibition activity. Three Csr/Rsm homologs are present in SBW25 and were named *rsmA1* (*PFLU4746*), *rsmA2* (*PFLU4324*) and *rsmE* (*PFLU4165*). The phenotypic effect of the Gac/Rsm pathway was investigated by creating deletion mutants for the response regulator *gacA* and the two Csr/Rsm homologs *rsmA1* and *rsmE*. Capsulation was completely abolished in a *gacA* deletion strain, confirming the transposon-mutagenesis results (Fig. 2b). Deletion of *rsmA1* or *rsmE* increased the production of capsulated cells, consistent with their typical inhibitory role in Gac/Rsm signalling pathways.

We postulated that variations in the relative concentration of free ribosomes and RsmA/E may determine translational output of a key positive regulator of capsule biosynthesis (Fig. 2c). While searching for such a regulator, attention turned to *PFLU3655*. The first gene of a putative operon (*PFLU3655-3657*) localized just upstream of the colanic acid biosynthetic operon, *PFLU3655* is annotated as a hypothetical protein carrying a two-component response regulator C-terminal domain (PFAM PF00486). *PFLU3655* is among the most highly up-regulated genes in 1B4 Cap^+^ cells and transposon insertions in its promoter were shown to abolish capsulation in 1B4 (ref. 9). A non-polar deletion of *PFLU3655* in 1B4 abolished capsule formation, while complementation of the mutant with an IPTG-inducible copy of *PFLU3655* on a low copy number plasmid (pME6032) restored capsulation (Fig. 3a). Over-expression of *PFLU3655* in SBW25 or 1B4 led to high capsulation levels in these strains, showing that PFLU3655 is a key positive regulator of colanic acid biosynthesis, the expression of which is sufficient for capsulation. Moreover, ectopic expression of PFLU3655 induces expression from its own promoter, as measured using the chromosomally-encoded P*pflu3655*-GFP translational fusion (Fig. 3b). This result shows that PFLU3655 expression can generate a positive feedback loop, a motif that can sustain bistable gene expression^20,21^.

**Figure 3:**
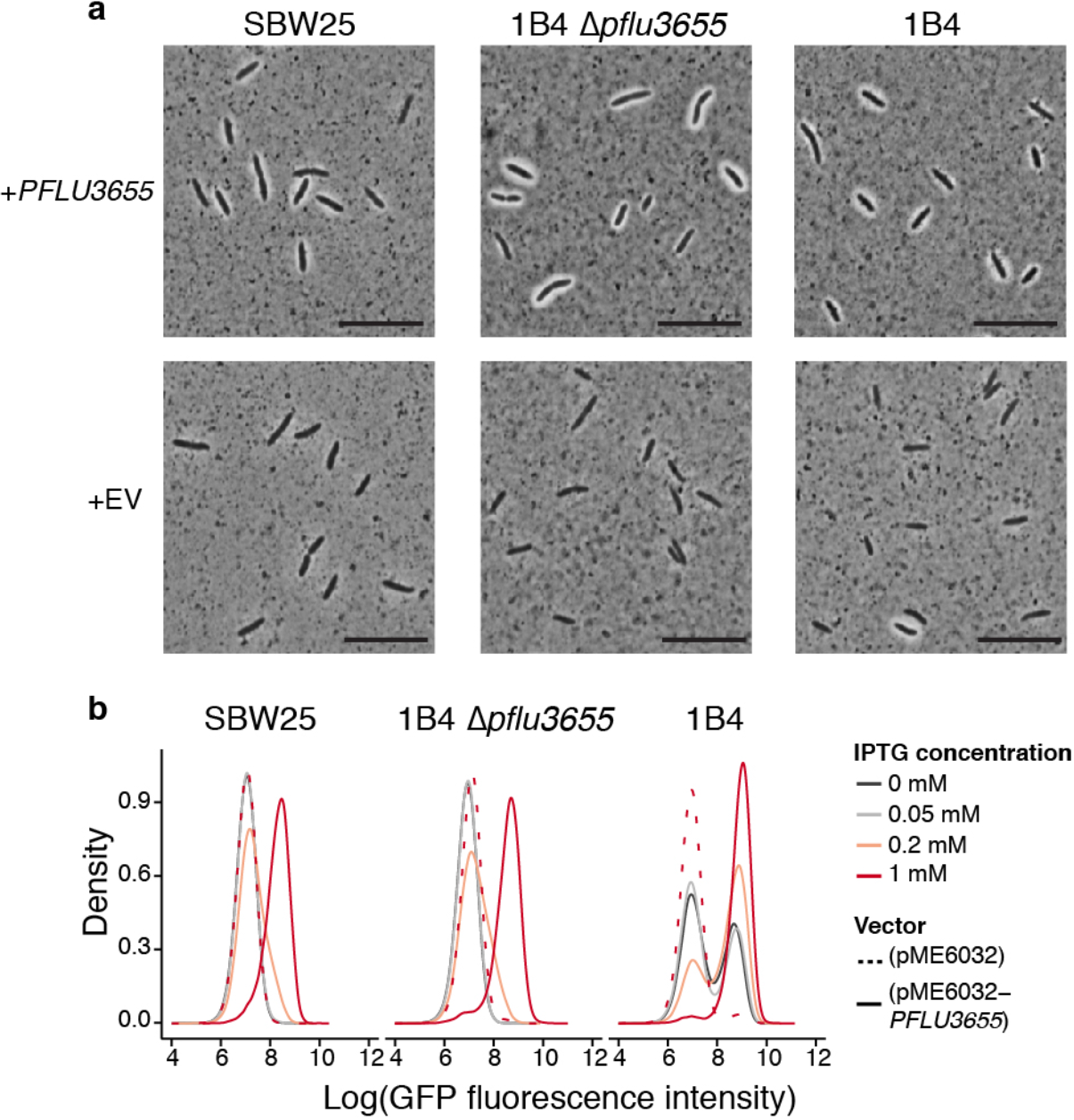
PFLU3655 is required for capsulation. **a**, Capsulation in SBW25, 1B4 ∆*pflu3655* and 1B4 strains carrying the pME6032-*pflu3655* plasmid or the empty vector (EV) after induction with 1mM IPTG. Phase contrast microscopy images of bacterial suspensions counter-stained with indian ink. White halos around cells indicate capsulation. Scale bar = 10 µm. **b**, PFLU3655 establishes a positive feedback loop. GFP fluorescence from the P*pflu3655*-GFP reporter in SBW25 (left), 1B4 ∆*pflu3655* (middle) or 1B4 (right) cells carrying the pME6032-*pflu3655* plasmid or empty vector and *pflu3655* expression was induced with IPTG at indicated concentration and fluorescence was measured by flow cytometry. Data are representative of 3 independent experiments (**a**, **b**).

Existence of a positive fedback loop does not however guarantee bistability, which often requires the additional presence of an ultrasensitive switch to convert small input deviations (typically, molecular noise) into large output differences^22^. When signalling components are present in large numbers, ultrasensitive responses and threshold effects can arise through molecular titration where a molecule (RNA or protein) is sequestered and inhibited by another protein^22–25^. This led to rocognition that titration by RsmA/E may be involved in phenotypic bistability.

Two putative Rsm binding sites are located in the promoter and 5’ region of the coding sequence of *PFLU3655* (Fig. 4a), indicating that PFLU3655 mRNA could be a direct target of RsmA/E. The P*pflu3655*-GFP capsulation reporter was used to test this hypothesis. This translational reporter contains ~500 nucleotides upstream of the PFLU3655 start codon with the first 39 coding nucleotides being fused in frame to GFP; the two putative RsmA/E binding sites are conserved in this synthetic construct. Using site-directed mutagenesis, nucleotides located in the putative Rsm binding sites were substituted and effects on GFP expression in the P*pflu3655*-GFP reporter strain determined (Fig. 4a,b). Both G-8A and A33T point mutations increased GFP production, albeit to different extents, consistent with the expected effects arising from reduction in binding of an inhibitor. Altering the putative RBS (GG-7AC) completely abolished GFP expression (Fig. 4b), which is similarly to be expected given the need for translation.

**Figure 4:**
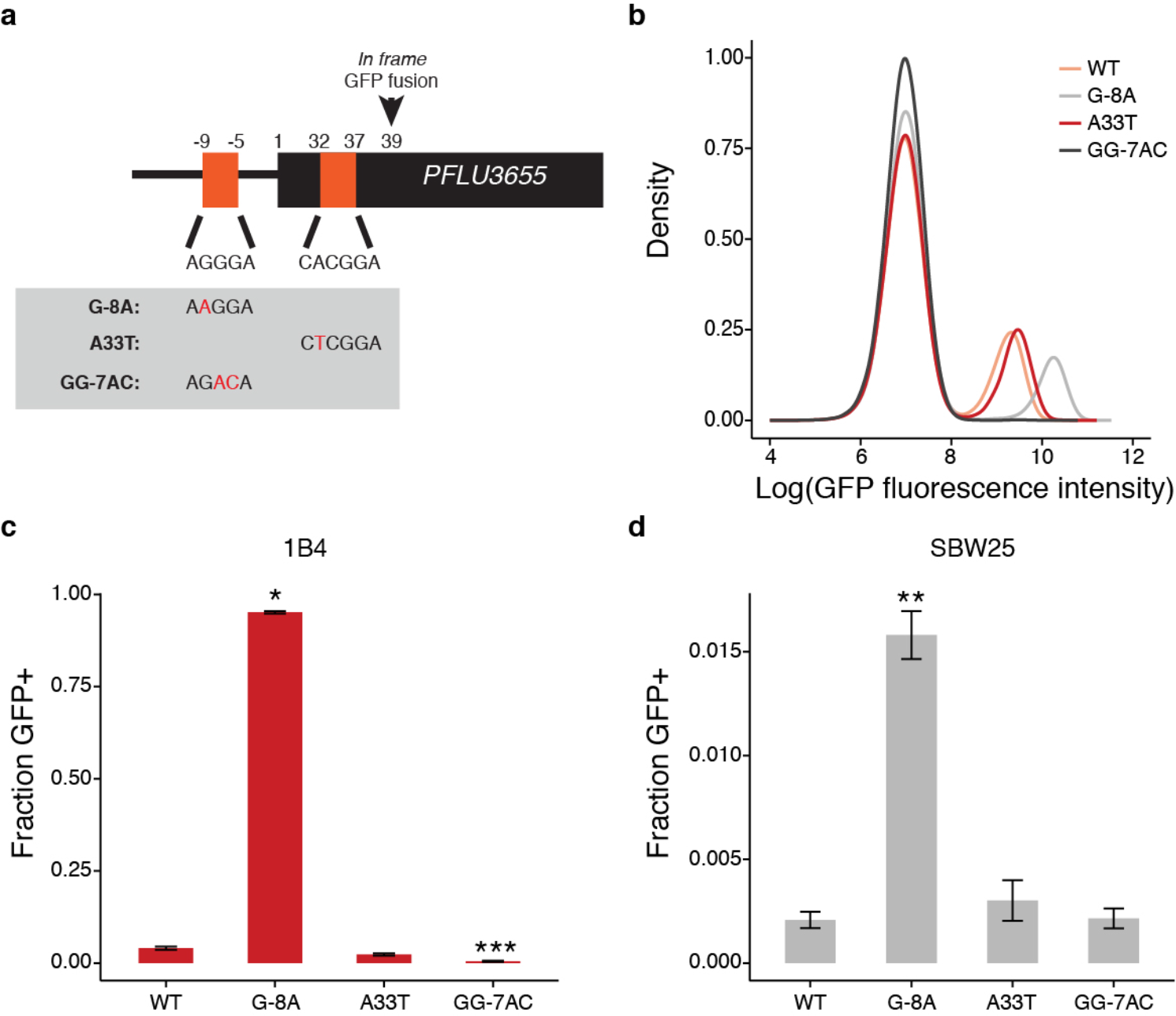
RsmA/E binding sites in *PFLU3655* control capsulation. **a**, Schematic diagram of *PFLU3655* region. Two putative RsmA/E binding sites (orange squares) are located in the promoter and 5’ region of the gene. Numbers indicate nucleotide positions relative to start codon (not to scale). Sequences of putative RsmA/E binding sites are shown, the putative ribosome-binding site (RBS) is underlined. Grey box: point mutations introduced in the different sequences by site-directed mutagenesis. **b**, Expression of the P*pflu3655*-GFP reporter carrying the different point mutations in the 1B4 background. GFP fluorescence was measured by flow cytometry. Data are representative of 3 independent experiments. **c-d**, Mutations in putative RsmA/E binding sites affect capsulation in 1B4 (**c**) and SBW25 (**d**). Individual point mutations were re-introduced into 1B4 and SBW25 carrying the wild-type P*pflu3655*-GFP reporter and the proportion of GFP positive cells in late exponential phase (OD_600nm_ = 1-2) was measured by flow cytometry. Means ± s.e.m. are shown, n = 9 (1B4) or n = 7 (SBW25). Data are pooled from 3 independent experiments. * *P* < 0.05, ** *P* < 0.01, *** *P* < 0.001, Kruskall-Wallis test with Dunn’s post-hoc correction, comparison to 1B4.

To determine effects on capsulation, the individual point mutations were re-introduced at the native locus in 1B4 and SBW25. The G-8A mutation, but not the A33T mutation, increased the proportion of capsulated cells in both strains (Fig. 4c, d). This difference mirrors the difference observed on expression of GFP and suggests that the increase in PFLU3655 translation mediated by the A33T mutation is not sufficient to increase the likelihood to jump-start the positive feedback loop, contrary to G-8A.

Together, these data support the model proposed earlier (Fig. 2c). If this model is correct, one would expect other RsmA/E targets to be over-expressed in *carB* mutants. A list of genes that were differentially expressed in a SBW25 *gacS* mutant^26^ was extracted and their expression levels were compared using the RNAseq dataset. On average, genes that were up-regulated in the *gacS* mutant were expressed at lower levels in 1B4 (both Cap^−^ and Cap^+^) than in SBW25 or 1A4 (Supplementary Figure 8). Genes that were down-regulated in *gacS* showed a slight bias towards higher expression in 1B4 Cap^+^ but this difference was not statistically significant. These results are consistent with the opposing effects of *gacS* inactivation (leading to constitutive activation of RsmA/E) and *carB*-dependent increase in ribosome concentration on RsmA/E targets.

### A qualitative mathematical model of post-transcriptional control of a positively regulated gene

We produced a mathematical model based on our experimental observations in order to describe the qualitative behaviour of the cells, notably bistability of the internal state, and to predict how the explored changes in the regulation pathways are likely to affect the fraction of capsulated cells.

An ordinary differential equation for the concentration of mRNA transcribed from *PFLU3655* formalizes the two hypotheses that transcription is positively regulated by the protein PFLU3655 concentration, and that ribosomes and the regulator RsmA/E compete for a binding site on the mRNA (Supplementary Note). The model, illustrated in Figure S1 from Supplementary Note, describes the dynamics of three mRNA pools: free, bound to ribosomes (thus translated) and bound to the regulator. Modifications in total ribosome concentration, in the rate of basal mRNA production, and in regulator binding efficiency can be included in this model in the form of parameter changes. By studying their effect on the system dynamics, and in particular its equilibria, the model can be used to check whether the hypotheses formulated previously on ribosome-mediated translational regulation are consistent with experimental observations.

Like other systems where post-transcriptional control is mediated by regulators acting as mRNA ’sponges’, molecular titration introduces nonlinearities that can give rise to bistability^22–25^. When the system is bistable, an ’OFF’ equilibrium, where the gene is not translated, coexists with an ’ON’ equilibrium, where the protein, thus the fluorescent reporter, are produced. The level of protein expression in this second equilibrium is set by saturation of the transcription rate with increasing protein concentration. These two stable equilibria are separated by a third unstable equilibrium – whose position depends on all the parameters of the system – which sets the protein concentration threshold for mRNA translation to overcome sequestration by the regulator. Assuming that the transition between the two alternative states takes place due to stochastic processes at the molecular level, extension of the basin of attraction of either stable equilibrium can be taken as a proxy of the probability of observing cells in the corresponding state.

As shown in Figure S2 from Supplementary Note, the model reproduces the qualitative modifications observed experimentally: the entry into a bistable regime when ribosome levels increase (hence the difference between SBW25 and 1B4 strains), and the increase both in the capsulation probability and in the protein levels when either a second, unregulated, source of mRNA production is added, or the binding affinity of the regulator is reduced. As a corollary of our model ribosome levels are expected to be heterogeneous in the population with capsulated cells having an increased average ribosome content with respect to non-capsulated cells. Indeed, RNAseq data indicate that ribosomal protein genes are expressed at higher levels in 1B4 Cap^+^ compared to 1B4 Cap^−^ (Fig. 1a).

### Consequences of ribosome heterogeneity on growth resumption in capsulated cells

When the quality and/or quantity of nutrients rises abruputly, differences in ribosome abundance can be significant for bacterial fitness^27^. Reaching higher ribosome concentrations – required for maximal growth rate after nutrient up-shift – is time-consuming and introduces a time-delay between environment change and future growth^28^. Cells with higher ribosome concentrations before nutrient up-shift might be considered as having been provisioned for rapid acclimation to the new conditions. If it is true that ribosomes promote growth resumption after nutrient up-shift, then capsulated cells should have an average growth advantage under these conditions. To test this prediction, 1B4 cells grown to late exponential phase (OD_600nm_~1) and cell suspensions enriched in Cap^−^ or Cap^+^ cells were used to inoculate fresh cultures. The initial growth rate after nutrient up-shift in batch cultures was positively correlated with the proportion of capsulated cells (Fig. 5a). Time-lapse microscopy on solid agar pads confirmed that colonies founded by GFP^+^ (capsulated) cells grew approximately 10% faster than those originating from GFP^−^ (noncapsulated) cells (Fig. 5b).

**Figure 5:**
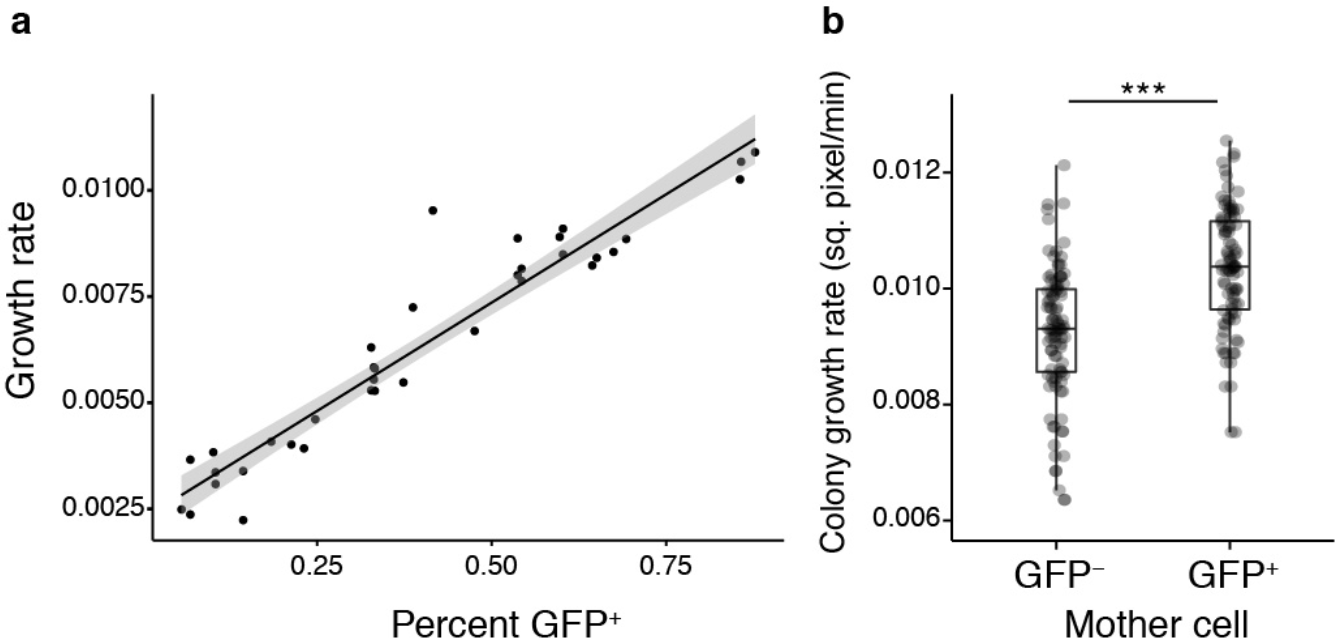
Capsulation and growth in 1B4. **a**, Initial growth rate after nutrient upshift is correlated to the proportion of capsulated cells in 1B4 populations. Data points are pooled from 2 independent experiments. n = 36, r^2^ = 0.91. Shaded area indicates 95% confidence interval. **b**, Growth rate of micro-colonies founded by Cap^−^ (GFP^−^) or Cap^+^ (GFP^+^) cells measured by time-lapse microscopy. n = 97 (GFP^−^) or n = 94 (GFP^+^). Data are pooled from 7 independent experiments. *** *P* < 0.001, two-tailed *t*-test

### Capsulation in SBW25

Identification of the mechanism promoting production of colanic acid capsules in the derived 1B4 switcher genotype raises questions as to the role and conditions for expression of colanic acid in ancestral SBW25. In bacteria, extracellular capsules are important for bacterial pathogenicity^29^ and are associated with broader environmental versatility^30^. Recent work has also demonstrated a role for colanic acid-like capsules in the positioning of cells at the surface of bacterial colonies, providing access to oxygen and a fitness advantage^31,32^.

In order to test if capsulation occurs in ancestral SBW25 colonies, we spot-inoculated bacterial suspensions on agar plates with incubation at 28°C for several days. Mucoid papillations were observed in the centre of colonies from 5 days post-inoculation and increased over time (Fig. 6a). These papillations were not observed in colonies derived from a colanic acid mutant (data not shown) and their appearance was delayed when 2mM uracil was added to agar plates (Fig. 6b). In SBW25, cells sampled from mucoid regions showed a high proportion of capsulated cells, approximately 60% of which expressed the P*pflu3655-*GFP reporter (Fig 6c). The fact that some capsulated cells do not express the GFP reporter in old colonies may result from remanence of capsules around cells that have stopped production, and/or from a possible PFLU3655-independent capsulation program. When streaked on new agar plates, no phenotypic difference was observed in colonies arising from mucoid versus non-mucoid cells (data not shown), indicating that mucoidy is not dependent on *de novo* mutation. These results indicate that capsulation in starved SBW25 colonies is also a consequence of bistable colanic acid production.

Given previous results with 1B4, we reasoned that capsulated SBW25 cells might also benefit from a growth advantage upon exposure to rich medium. Cells were collected from 7-day old colonies and enriched in capsulated or non-capsulated cells by gentle centrifugation. When transfered to fresh batch cultures, cell suspensions enriched in capsulated cells showed a faster initial growth rate (Supplementary Figure 9). To directly measure the fitness effect of capsulation during growth resumption, cellular suspensions from colonies of SBW25 or its isogenic variant carrying a neutral *lacZ* marker^33^ were collected. SBW25 Cap^−^ cells were mixed with SBW25-*lacZ* Cap^+^ cells, and *vice versa*, to initiate competition experiments. A significant fitness advantage of capsulated cells was detected after 2h or 4h growth in KB medium (Fig. 6d). Finally, time-lapse microscopy experiments revealed that micro-colonies founded by GFP^+^ cells display an initial growth rate significantly higher than colonies arising from GFP^−^ cells (Fig 6e).

**Figure 6:**
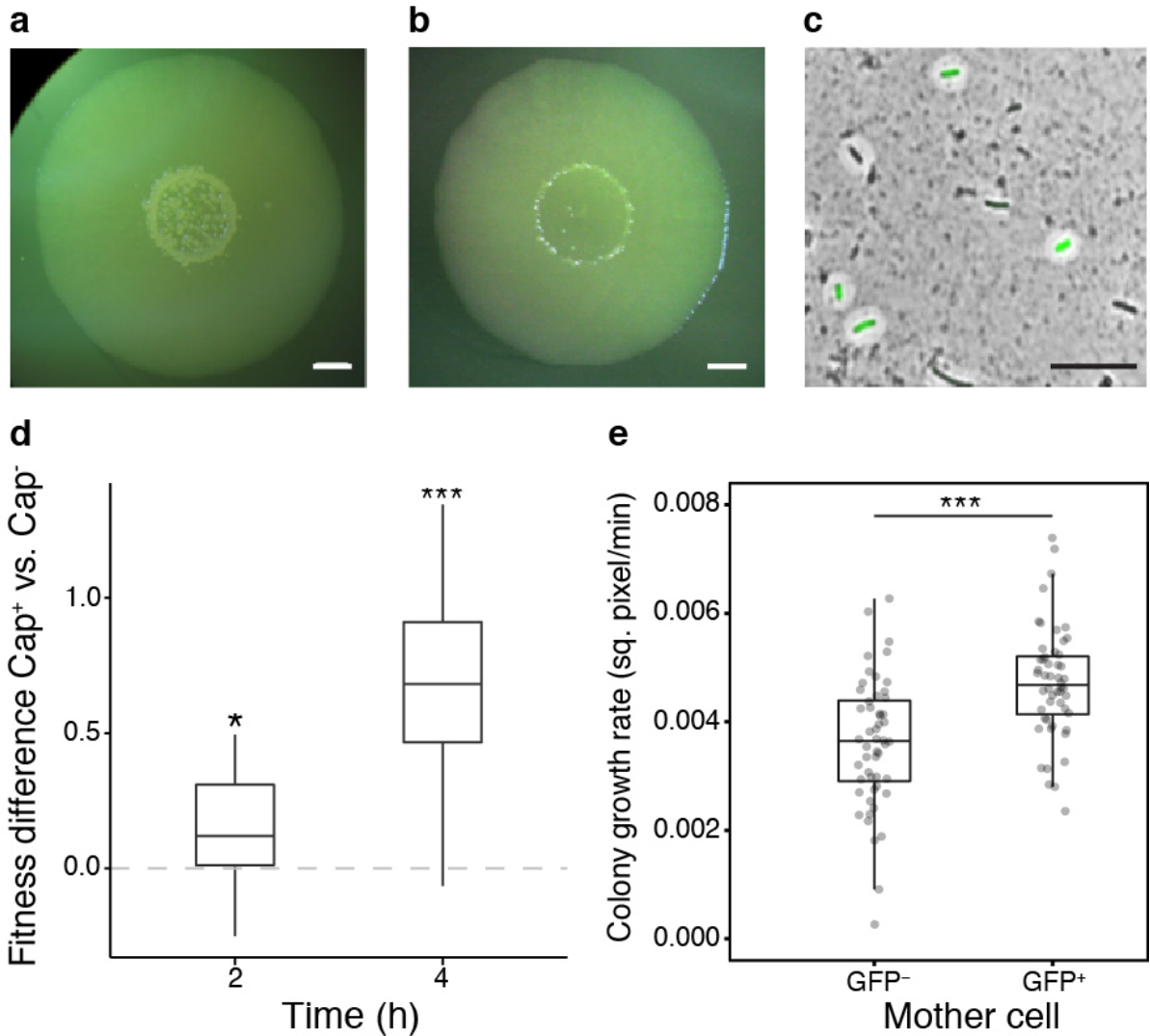
Capsulation and growth in SBW25. **a-b**, SBW25 colony grown on KB agar plate (**a**) or KB agar plate supplemented with 2 mM uracil (**b**) for 7 days. Scale bar = 2 mm. **c**, SBW25 cells carrying the Tn7-P*pflu3655*-GFP reporter and sampled for a 7-day old colony were counter-stained with indian ink to detect the presence of colanic acid capsules. A GFP image are overlaid with the phase contrast image. Scale bar = 10 µm. Images are representative of at least 3 independent experiments (**a-c**). **d**, Competitive fitness difference between SBW25 Cap^−^ and Cap^+^ cells. Boxplot of the differences in Malthusian parameters between cultures enriched in Cap^+^ vs. Cap^−^ cells are shown, n = 12. Data are pooled from 2 independent experiments. * *P* < 0.05,*** *P* < 0.001, comparison to 0 with two-tailed *t*-test. **e**, Initial growth rate of micro-colonies founded by GFP^−^ or GFP^+^ cells measured by time-lapse microscopy. n = 65. Data are pooled from 4 independent experiments. *** *P* <0.001, Wilcoxon test.

To test if capsulation status in SBW25 is also associated with higher average ribosome content, we measured expression of the ribosomal protein gene *rpsL* in capsulated and noncapsulated cells originating from old SBW25 colonies by RT-qPCR. We found an expression ratio of 1.67 (+/− 0.35 s.d., n = 5, two-tailed *t*-test, *P* = 0.012) in Cap^+^ vs. Cap^−^ cells, suggesting that SBW25 capsulated cells contain on average more ribosomes than their non-capsulated counter-parts.

Overall, the results from experiments carried out in SBW25 are consistent with the capsulation model proposed for 1B4. During ‘long-term’ (*i.e*., 7 days) starvation, nutrient limitation (possibly triggered by a reduction in flux through the pyrimidine pathway) causes a shift towards increased ribosome content. Cells with higher ribosome levels have an enhanced chance of flipping to the capsulated state where growth remains slow and cells enter a semi-quiescent state. Even though giving up competition for nutrients in stationary phase, these cells then stand primed for rapid growth upon nutrient upshift.

## Discussion

Studies of adaptive phenotypes derived from selection experiments are by nature multifaceted, but two parallel lines of inquiry are of particular importance. The first concerns the nature of the adaptive phenotype, including its selective and molecular causes. The second concerns the ecological significance of the traits affected by adaptive evolution, prior to the occurrence of the adaptive mutation(s). A satisfactory answer to the first requires understanding of the latter.

Early characterisation of the 1B4 switching genotype^8^ showed the behaviour to be a consequence of a single non-synonymous nucleotide substitution in carbamoyl phosphate synthase *carB*. In and of itself discovery of the causal switch-generating mutation shed no light on the adaptive phenotype. Understanding began to emerge only upon recognition that the *carB* mutation had altered the activation threshold of a pre-existing switch^9^. Here we have substantially extended understanding of the mechanistic bases of phenoytpic switching in the derived 1B4 genotype.

Experiments examining function of the bistable behaviour in a *galU* mutant of the 1B4 switcher showed that previous conclusions concerning the central role of UTP or related molecules required revision. While pyrimidine biosynthesis and the UTP decision point are important components of the pathways leading to capsulation, the mechanism of switching resides elsewhere. Data presented here have led to formulation of a compelling new model of the switch. Central to the proposed model (Fig. 2c) is competition between RsmA/E and ribosomes for *PFLU3655* mRNA, a positive regulator of capsulation. That translation initiation and/or efficiency may affect signaling through the GacAS two-component system was previously suggested in two independent studies^34,35^.

Importantly, in our model, titration of *PFLU3655* mRNA by RsmA/E has the potential to generate bistability. An increase in ribosome production resulting from the *carB* mutation leads the system to a bistable regime. Small fluctuations in ribosome and/or in RsmA/E activities beyond an activation threshold initiate a positive feedback loop leading to capsulation. The position of the threshold and the consequent probability of switching are affected by even small changes in parameters of the system. The prevalence of inhibitory interactions in signaling networks offers a common route for the evolution (or evolutionary tuning) of bistable switches through relatively minor changes to the basal level or the interaction affinity of threshold-defining components^6,23,36^. Bimodal expression of GacAS-regulated genes has been observed in *Pseudomonas aeruginosa*^37^ and the relevance of ribosome-mediated heterogeneity for numerous two-component signaling pathways involving RNA-binding proteins deserves further attention. Such regulatory strategies would enable bacterial cells to couple their internal metabolic and physiological status to multiple signaling outputs. Heterogeneity in persister resuscitation in *E. coli* was recently shown to correlate with ribosome content^38^ and provides a further example of the association between ribosomes and phenotypic heterogeneity.

A remaining open question is the mechanistic link between pyrimidine starvation and ribosome biosynthesis. While the stringent response is believed to tune ribosome production to cellular needs^39^, a phenomenon of ribosome over-capacity – bearing similarity to what we describe as ‘ribosome provisioning’ – was previously reported in slow growing bacteria^40–42^ and occurs concomitantly with a reduction in the rate of translation (or accumulation of inactive ribosomes). RelA-dependent production of ppGpp was shown to be necessary for ribosome accumulation under nitrogen starvation^42^, but a *relA spoT* mutant of genotype 1B4 was unaffected in its capacity to switch (data not shown). It is possible that in 1B4 an imbalance in the nucleotide pool – a factor known to influence ribosome production in *E. coli*^43^ – may directly alter ribosome production.

*In vitro* selection experiments can shed light on hitherto unrecognized aspects of bacterial physiology^44–49^. Understanding the evolutionary origin of the switch between cells with differing capsulation states requires understanding of the function and ecological significance of the switch in the ancestral genotype. Previous work showed that switching to the capsulated type was accompanied by a reduction in growth rate and that the probability of switching to the capsulated state was more likely in starved cells (particularly in cells starved of pyrimidines). This was understood as a mechanism that allowed cells entering starvation conditions to hedge their bets in the face of uncertainty surrounding the future state of the environment^9^. Discovery that capsulated cells, despite slow growth, are replete in ribosomes was perplexing, but caused attention to focus on exit from the semi-quiescent state. Just as cells entering a slow growing phase stand to be out-competed by conspecific types that remain in the active growth phase should the environment unexpectedly return to one conducive for growth, cells exiting from a slow growth state stand to be outcompeted by types that are already actively growing unless they can rapidly resume “life in the fast lane”. This phenomenon that we refer to as ‘ribosome provisioning’ has parallels with recent reports of ribosome dynamics in exponentially growing cells^27,50,51^. In environments where resource availability fluctuates, it appears that the control of ribosome biosynthesis is subject to a trade-off between maximising growth rate during nutrient limitation and growth resumption upon nutrient up-shift. This trade-off may be solved at the single-cell level, where heterogeneity in ribosome activity may contribute to optimize long-term geometric mean fitness.

Although the subject of little attention in the microbiological world (but see refs. 27, 51, 52) ideas concerning provisioning of future generations with resources sufficient to aid their establishment is a component of life-history evolution theory^53^. It has been particularly well developed in the context of seed dormancy and the evolution of post-germination traits^54^. It is not difficult to conceive that bacterial cells entering a slow or non-growing state, such as persisters, or the capsulated cells of SBW25, will through evolutionary time, experience selection for mechanisms that facilitate rapid re-entry to active growth. Our data here is suggestive of such an evolutionary response.

## Material and methods

### Bacterial strains and growth conditions

Bacterial strains used in this study are listed in Supplementary Table 1. *Pseudomonas fluorescens* strains were cultivated in King’s Medium B (KB; ref. 55) at 28°C. *Escherichia coli* DH5α *λpir* was used for cloning and was grown on Lysogeny Broth at 37°C. Bacteria were plated on their respective growth media containing 1.5% agar. Antibiotics were used at the following concentrations: ampicillin (50-100 μg mL^−1^), gentamicin (10 μg mL^−1^), tetracycline (10 μg mL^−^1), kanamycin (25 or 50 nitrofuorantoin (100 μg mL^−^1 for *E. coli* or *P. fluorescens*, respectively) and μg mL^−^1). Uracil (Sigma-Aldrich) was added to culture medium at 2 mM final concentration when indicated. For competition experiments, 5-bromo-4-chloro-3-indolyl-β-d-galactopyranoside (X-gal) was used at a concentration of 60 mg L^−1^ in agar plates.

For capsulation assays, pre-cultures were inoculated from pre-calibrated dilutions of frozen glycerol aliquots in order to reach an OD600nm of 0.3-0.5 after overnight culture.

For colony assays with SBW25, 5 μl of cell suspensions were spot-inoculated on KB agar plates and incubated for 7 days at 28°C. Cells from the center of these colonies were resuspended in PBS or Ringer’s solution for growth and competition assays and time-lapse microscopy, or in RNAlater solution (Invitrogen) for RT-qPCR.

### Molecular techniques

Oligonucleotides and plasmids used in this study are listed in Supplementary Tables 2 and 3, respectively. Standard molecular biology techniques were used for DNA manipulations^56^. DNA fragments used to generate promoter fusions and gene deletion constructs were prepared by splicing by overhang extension polymerase chain reaction (SOE-PCR; ref. 57). All DNA fragments generated by SOE-PCR were first cloned into the pGEM-T easy vector (Promega) and their fidelity was verified by Sanger sequencing (Macrogen, Seoul). Plasmids were introduced into *P. fluorescens* by tri-parental conjugations with the helper plasmid pRK2013 (ref. 58), carrying the *tra* and *mob* genes required for conjugation. Tn7-based plasmids were mobilized into recipient strains with the additional helper plasmid pUX-BF13 (ref. 59).

To generate deletion mutants (*rrn* operons, *gacA, rsmA, rsmE* and *pflu3655*), regions flanking the genes or operons of interest were amplified from SBW25 genomic DNA and assembled by SOE-PCR. Deletion cassettes were inserted into the pUIC3 plasmid^60^ as *Spe*I fragments and mutants were obtained following the two-step allelic exchange protocol described previously^33^. Deletion mutants were checked by PCR. To check *rrn* copy number after *rrn* deletions, quantitative PCR was performed using a protocol described previously^61^.

For complementation and over-expression studies, *PFLU3655* was amplified and cloned into pME6032 (ref. 62) as an *EcoR*I/*Xho*I restriction fragment, downstream of the P*tac* promoter.

To generate the Tn7-P*rrnB*-GFP reporter, a ~600bp fragment upstream of the *rrnB* operon (*PFLUr7-11*) was amplified from SBW25 genomic DNA and fused by SOE-PCR to *gfpmut3* sequence containing the T0 terminator previously amplified with oPR152/FluomarkerP2 from the miniTn7(Gm)-P*rrnB1*-*gfpmut3* plasmid^63^. The resulting fragment was cloned into pUC18R6K-mini-Tn7T-Gm (ref. 64) as a *Spe*I restriction fragment.

Site-directed mutagenesis of putative RsmA/E binding sites in pGEMT easy-P*pflu3655*-GFP plasmid was performed using the Quick Change mutagenesis kit (Stratagene) according to manufacturer’s instructions. Mutagenized fragments were then cloned into pUC18R6K-miniTn7T-Gm (ref. 64) as a *Spe*I restriction fragment. In order to re-engineer point mutations in RsmA/E into the *Pseudomonas fluorescens* genome, a 1.5kb fragment spanning equal length on each side of the target sites was amplified and cloned into pGEM-T easy vector. Site-directed mutagenesis was performed on this plasmid as described above, and the resulting DNA fragments were cloned in pUIC3 as *Spe*I restriction fragments and introduced into the *P. fluorescens* genome *via* the two-step allelic exchange protocol.

### RNA extractions and RT-qPCR

For quantification of RNA concentration in bacterial cultures, cells were harvested from 1 ml of cultures at OD_600nm_ of 0.5-0.6 and resuspended in 200 μl of RNAlater solution (Invitrogen). For total RNA quantification, we followed the method described by ref. 65, except that, before processing, cells cultures were resuspended in RNAlater (Invitrogen) instead of being fast frozen on dry ice. To normalise total RNA concentrations, the relationship between cell density and OD_600nm_ was established for each strain by counting cells with a hemocytometer in 5 independent cultures of similar OD to those used for RNA extractions. For *rrn* double mutants, no significant difference in the cell/OD_600nm_ ratio was found when compared to 1B4, so RNA quantities were normalised with OD_600nm_ values.

Reverse-transcription quantitative PCR was performed as described previously^61^, using *gyrA* as an internal control. Oligonucleotide primers used for RT-qPCR are listed in Supplementary Table 2.

### Capsulation and gene expression assays

For capsulation tests, cells were grown from standardized glycerol aliquots stored at −80°C. Aliquots were diluted in KB and pre-cultures were grown overnight in order to reach an OD_600nm_ of 0.3-0.5 in the morning. For gene expression studies, overnight pre-cultures were grown to saturation. In both cases, pre-cultures were diluted to OD_600nm_of 0.05 in KB and incubated at 28°C. IPTG was added to a final concentration of 0.1-1mM when indicated. Samples were taken at different time points for flow cytometry and OD measurements. GFP fluorescence in bacterial populations was measured with a BD FACS Canto II flow cytometer. Cell suspensions were diluted to a density of ~ 10^5^ cells ml^−1^ in filter-sterilised PBS and at least 20,000 cells were analysed by flow cytometry. Cellular debris were filtered using side-scatter channel (SSC-H/SSC-W). GFP fluorescence was detected with a 488 nm laser with 530/30 bandwidth filter. Laser intensity was set to 600V, except for P*rrnB*-GFP analyses where intensity was lowered to 300V. Flow cytometry data files were analysed in R (ref. 66) using the ‘flowCore’ package^67^. For capsulation experiments, the relative sizes of GFP positive and negative sub-populations were measured after manual thresholding of GFP intensity, following bi-exponential transformation of the FITC-H signal. The distributions of expression intensities in Figures 3b. and 4b. were smoothed in R using the ‘KernSmooth’ package^68^.

### Growth curves in microplate reader

Overnight precultures were adjusted to OD_600nm_ of 0.05 and 200 μl KB cultures were grown in 96-well plates. Cultures were incubated in a Synergy 2 microplate reader (Biotek) for at least 24h at 28°C with constant shaking and OD_600nm_ was read every 5 minutes.

To measure growth rates of cultures enriched in capsulated or non-capsulated cells, cells were harvested from late-exponential phase (OD_600nm_ ~ 1; 1B4) or 7-day old colonies (SBW25) and centrifuged (1 min, 3000 rpm). Supernatants and pellets were collected, representing sub-populations enriched in capsulated and non-capsulated cells, respectively. Cell suspension density was adjusted to OD_600nm_ of 0.05 in KB to start growth curves. Initial growth rates were calculated by performing a linear regression on the logarithm of the measure OD values during the first 2h of growth.

### Microscopy

Microscopy experiments were performed with an Olympus BX61 upright microscope equipped with an F-View II monochrome camera, a motorized stage and a temperature-controlled chamber set at 28°C. Devices were operated by the Cell^P or CellSens softwares (Olympus). Phase-contrast images were acquired with an oil-immersion 100x/N.A. 1.30 objective. GFP fluoresecence images were acquired with the following filter set: excitation (460-480 nm), emission (495-540 nm) and dichroic mirror (DM485).

Indian ink staining of capsulated cells was performed as described previously^9^. To determine cell size, exponentially growing bacteria were diluted 1:10 in KB and transferred on 1% agarose-KB gel pads. Cell sizes were determined from phase-contrast images with MicrobeJ (ref. 69). For time-lapse microscopy, bacteria were harvested from late exponential phase (1B4, OD600nm ~ 1-2) or from 7 day-old colonies (SBW25), diluted 1:1000 in KB and 2μl of the resulting suspension was immediately transferred on a gel pad (1% agarose KB) located on a glass slide within an adhesive frame (GeneFrame, Thermo-Fisher). When dry, a cross-section of the pad was removed with a razor blade in order to allow gas exchanges to occur; the preparation was sealed with a glass cover-slip and transferred to the microscope incubation chamber pre-heated to 28°C. One GFP image was taken before starting the experiment in order to determine the capsulation status of each cell and phase-contrast images were then recorded every 10 minutes.

Phase contrast images were segmented with Fiji (ref. 70) and individual colony areas were extracted. For each colony analysed, the capsulation status of the founding cell was determined manually based on GFP signal. The logarithm of the growth rate of individual 1B4 colonies was then fitted using a linear regression. For SBW25, segmented linear regressions were found to better fit the data and the slope of the first line was reported.

### RNAseq analyses

RNAseq data were published previously^9^. KEGG orthology terms for the SBW25 genome were downloaded from the KEGG Orthology database (www.genome.jp; accessed in April 2016). KEGG enrichment statistics were computed with a hypergeometric test and were performed separately for up- and down-regulated genes. Data from the transcriptome analysis of SBW25 *gacS* mutant were retrieved from Supplementary Table 3 from ref. 26.

### Competition experiments

SBW25 or SBW25-*lacZ* cell suspensions were spot-inoculated on separate KB agar plates and grown for 7 days. Cells from the centre of colonies were harvested, resuspended in Ringer’s solution and gently centrifuged to enrich suspensions in capsulated or non-capsulated cells. Capsulated SBW25 cells were mixed with non-capsulated SBW-*lacZ* cells, and vice-versa. Mixed suspensions were diluted 1:100 in KB medium and grown at 28°C with orbital shaking for 4h. Appropriate dilutions of the cultures were plated on KB + X-gal plates at 0h, 2h and 4h post-inoculation in order to measure the ratio of white-blue colonies. The difference in Malthusian parameters^71^ was used as a measure of relative fitness.

### Statistical analyses

All statistical analyses were performed in R (ref. 66). Parametric data were analysed with a two-tailed Welch’s *t*-test. Non-parametric data were analysed with a Wilcoxon test (2 samples) or with a Kruskall-Wallis test with Dunn’s post-hoc correction (multiple comparisons). In Figure 5a, r^2^ indicates the adjusted R-squared value calculated from the linear regression. All *P* values are provided in Supplementary Table 5. All measurements were performed on distinct samples. The number of replicates indicate biological replicates, consisting of independent bacterial cultures or individual cells/micro-colonies (Figures 5b, 6e and Supplementary Figure 4).

### Data availability

Data and material are available from corresponding authors upon reasonable request.

## Acknowledgements

The authors thank Jenna Gallie and Camille De Almeida for discussion, Xue-Xian Zhang for the generous gift of pUIC3-Δ*gacA* deletion plasmid, Heather Hendrickson and Peter Lind for help with microscopy and flow cytometry, respectively, and Elena Denisenko for assistance with KEGG enrichment analyses. This work was supported in part by the Marsden Fund Council and a James Cook Research Fellowship from government funding administered by the Royal Society of New Zealand. SDM has received support under the program « Investissements d’Avenir » launched by the French Government and implemented by ANR with the references ANR-10-LABX-54 MEMOLIFE and ANR-10-IDEX-0001-02 PSL* Research University.

## Author contributions

PR and PBR designed the study, PR and GCF performed experiments and analysed the data, SdM designed and analysed the mathematical model, PR and PBR wrote the paper with contributions from all authors.

